# Leveraging defense system modularity to discover anti-phage systems

**DOI:** 10.64898/2025.12.08.692795

**Authors:** Yanqiu Liu, Keyi Tan, Zhenhao Han, Yu Chen, Kangyi Xiao, Pian Luo, RuiPeng Fan, Fuming Liang, Rafael Pinilla-Redondo, Wenyuan Han

## Abstract

Bacterial exposure to constant phage attack drives rapid diversification of anti-phage defense systems, often through the exchange of modular defensive domains. Here, we leverage this modularity signature to identify new defense systems by systematically searching for operons encoding defensive domains in non-canonical configurations.

We identified 214,164 candidate defense operons in *E. coli* genomes, representing a ∼2.2-fold expansion over defense systems detected by DefenseFinder. Experimental testing of 9 candidates validated 6 with anti-phage activity. These include DarTG and ietAS system variants that have acquired helicase modules, and a Gabija system in which a MazF-like protein replaces GajA, implying novel anti-phage mechanisms.

We also identified a new clade of Pycsar and show that it synergizes with type IV Thoeris to broaden phage protection. Our findings demonstrate that mining modular defensive domains provides a powerful strategy to predict and characterize new anti-phage systems, expanding the known repertoire of bacterial immunity.

**Teaser:** Modular domains of anti-phage defense systems are harnessed to predict novel defense systems

## Introduction

Bacteria are under constant threat of infection by phages (*1*). The resulting selective pressure has driven the evolution of a diverse and rapidly expanding arsenal of anti-phage defense systems (*2–4*). These systems can directly target and destroy phage genomes, such as restriction-modification (RM) (*5*) and CRISPR-Cas systems (*6*), thereby protecting the infected cell. Others interfere with essential host processes to inhibit phage replication (*7*). In these cases, although the infected cell dies or enters a dormancy state, uninfected cells in the community will survive, resulting in population-level immunity.

A previous quantitative analysis of the prokaryotic anti-phage arsenal revealed that bacterial genomes typically encode 3-8 defense systems (*8*). Since then, the anti-phage arsenal has continued to expand (*9–17*), indicating that novel anti-phage systems remain to be discovered. The discovery of many defense systems has relied on the “guilt-by-association” principle (*9, 18, 19*), i.e., defense systems tend to co-localize in genomic “defense islands”, enabling prediction of candidate defense systems based on proximity to known systems (*20*). In addition, defense systems are enriched in certain genomic locations of mobile genetic elements (MGEs), such as phage satellites (*10*), prophages (*15*), integrons (*12, 13*), and anti-phage testing of genes in these loci has further expanded the bacterial immune repertoire. Finally, functional screening of metagenomic fragments also represents a powerful method for identifying defense systems (*11, 14*).

A striking property of defense systems is their modular organization. Many systems share common functional domains that appear in different architectures (*2, 3*). For example, Sir2 domains act as NAD^+^-consuming effectors in a wide variety of systems (*9, 18, 19, 21–23*). This observation highlights that frequent module exchange facilitates the diversification of the anti-phage arsenal, suggesting that scanning for novel combinations of existing defensive modules can reveal previously uncharacterized defense systems (*21, 24–32*). Inspired by the modular nature of anti-phage systems, we developed a defense system prediction approach that systematically detects operons containing non-canonical combinations of defensive domains. Our results suggest that the candidate defense operons may outnumber complete defense systems detected by DefenseFinder (*8*) by ∼ 2.2-fold. From these candidates, we validated 6 operons as functional anti-phage systems. We also uncovered the synergistic effect of type IV Thoeris and a new clade of Pycsar systems. Together, our findings highlight the power of mining operons encoding defensive domains for the discovery of defense systems and provide insights into the diversification of the anti-phage arsenal during the evolutionary arms-race with phages.

## Results

### Prediction of candidate defense systems by domain association

Diversification through the exchange of conserved functional domains is a hallmark of bacterial anti-phage systems (*2*). We hypothesized that searching for operons encoding defensive domains, particularly in non-canonical combinations, could be an effective method for identifying uncharacterized anti-phage systems. DefenseFinder uses Hidden Markov Models (HMMs) of proteins involved in defense to detect known anti-phage systems (*8*), and therefore, also generates a list of proteins sharing homologous domains with the defense-associated proteins. We reasoned that these proteins carrying defensive domains represent a valuable resource for identifying novel defense systems.

To explore this idea, we scanned 4,243 *E. coli* genomes with DefenseFinder, which detected 17,988 complete defense systems (Table S1, Fig. S1A) and 551,238 proteins containing defensive domains (Fig. 1A, Fig. S1B). Of these, 356,990 proteins were not part of known systems and were used as seeds to search for operons encoding them. We removed the seeds not encoded within an operon, thereby excluding single-gene defense systems from further analysis. Most operons were automatically annotated as known systems based on their seed domain, although such labels do not necessarily reflect true system identity. For example, the Kongming system (*23*) was annotated as Thoeris due to the presence of a ThsA-like Sir2 domain KomC. A few operons were annotated with the names of the HMM domains built into DefenseFinder, for example, WYL and DEDDh. This approach yielded 214,164 unique operons (Table S2), spanning 98 distinct types of defense system. However, an analysis of their distribution across the *E. coli* genomes revealed that 21 operons, including Gao_Qat, CBASS and Dpd, are present in almost every strain (Fig. S1C), indicating that our set also included housekeeping proteins likely due to their homology to defense-associated proteins. To enrich for bona fide defense candidates, we applied the defense “guilt-by-association” principle and retained only those located within 50 kb of at least one defense system (Fig. 1A). This reduced the number of seeds to 71,253 and resulted in 39,848 unique candidate defense operons (CDOs) (Table S3), encompassing 82 defense system types. This represents a ∼2.2-fold increase over the number of complete defense systems detected by DefenseFinder (Table S1), consistent with the idea that a substantial fraction of the anti-phage repertoire in *E. coli* remains unexplored.

**Fig. 1.**
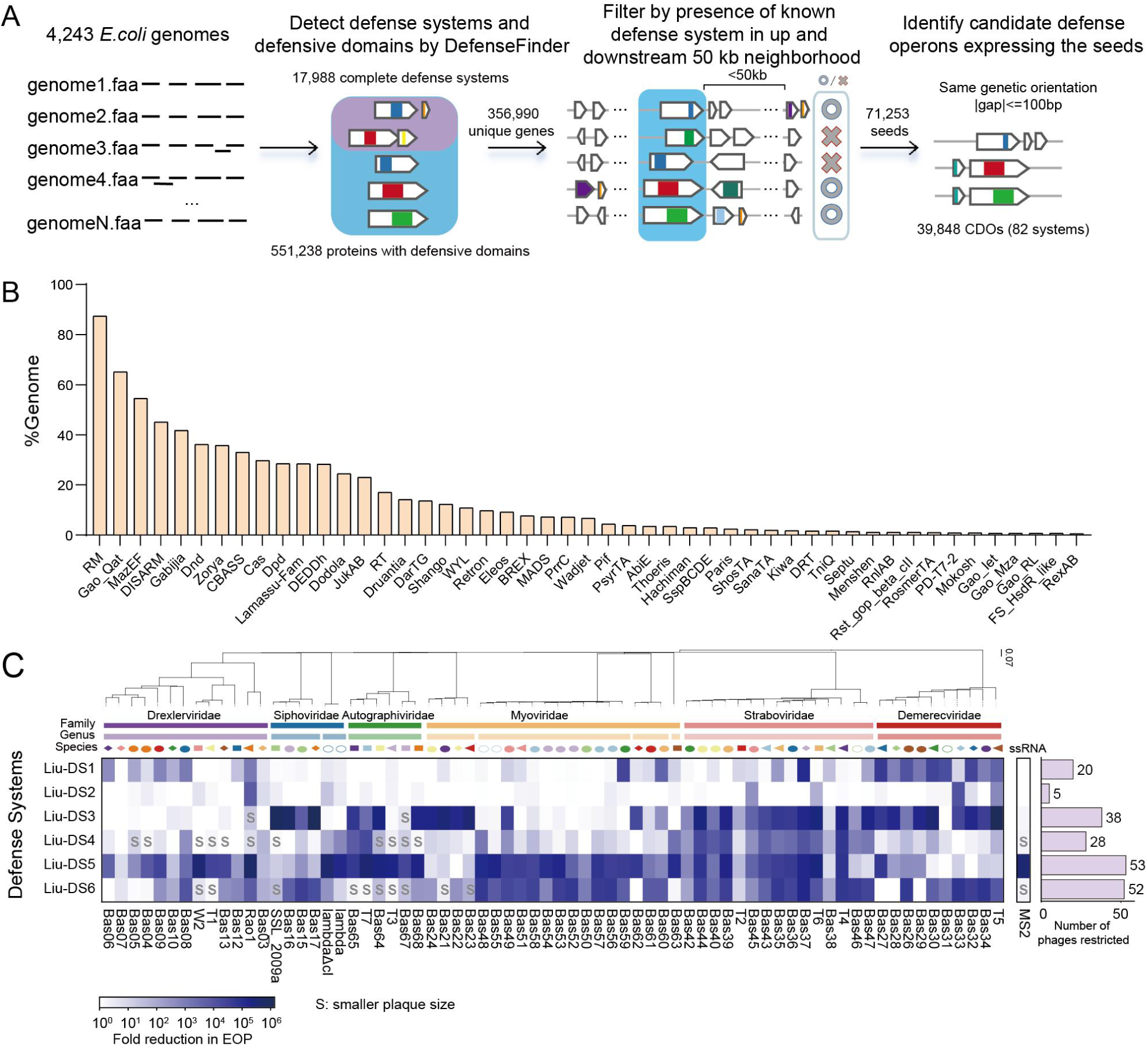
Prediction of candidate defense operons (CDOs) in *E. coli* genomes and functional validation. (A) Computational pipeline to identify CDOs in 4243 *E. coli* genomes. CDOs are defined based on the coding of a defensive domain and the presence of at least a known defense system within upstream and downstream 50 kb from the defensive domain. (B) Frequency of CDOs in *E. coli* genomes. Only CDOs present in >0.5% *E. coli* genomes are shown. (C) Defense heatmap of the anti-phage activity of validated defense systems, showing reduction in efficiency of plating (EOP) by blue squares and smaller plaque size indicated by the letter “s”. The decrease fold of EOP was calculated by 3 independent experiments. Phage restriction was defined by log(EOP reduction fold) > 1.5, and the numbers of restricted phages were counted.

We then analyzed the distribution of CDOs across *E. coli* genomes, according to their annotations. Restriction-modification (RM) CDOs were the most abundant, with 9,173 operons present in ∼90% *E. coli* genomes (Fig. 1B), compared to 6,556 complete RM systems in ∼70% genomes. Other abundant CDOs include Gao-Qat (65%, n = 3,131), MazEF (55%, n = 2,785), DISARM (45%, n = 2,182) and Gabija (42%, n = 2,038) (Fig. 1B). Inspection of their operon architectures showed that our pipeline identifies 1) complete systems with altered gene order, 2) defensive modules associated with non-defensive proteins, 3) mixed components from distinct systems, and 4) operons containing two different defense systems (Fig. S2). The combination of 2 defense systems may indicate synergistic effects, while new combinations with either defensive or non-defensive domains point to potentially novel anti-phage systems. However, we cannot exclude the possibility that some CDOs are not defense systems. For example, our analysis detected a twin-arginine translocation (Tat) system (Fig. S2), likely because the TatD nuclease is homologous to the QatD nuclease of Gao-Qat system.

To select candidate operons for functional testing, the 39,848 CDOs were clustered at 90% amino acid sequence identity, yielding 3,919 representatives (Table S4). RM CDOs (n = 1,343) and Cas CDOs (n = 308) were excluded because these systems have been studied extensively. The remaining CDOs were examined manually, removing those encoding complete defense systems with a rearranged gene order or including a transposase. From this curated set, we selected 9 CDOs candidates (Table S5) for functional characterization. These candidates were cloned into *E. coli* MG1655 and challenged with a broad panel of coliphages. This revealed that 6 out of the 9 tested candidates (hereafter designated Liu-DS1-6) conferred robust defense against at least one phage (Fig. 1C). One of these, Liu-DS5, contains an AVAST-associated metallo-β-lactamase (MBL)-like protein (Fig. S3A) and corresponds to HamMAB, recently described by Tuck et al. (*33*). Consistent with that study, Liu-DS5 restricted a wide range of phages (53/71), including an ssRNA phage (Fig. 1C), and provided population-level protection (Fig. S3B). Its immunity required the active MBL, protease, and helicase domains (Fig. S3C). Together, these results demonstrate that our domain-centric strategy successfully identifies bona fide anti-phage systems, motivating characterization of the additional systems described below.

### Potential nuclease expands the Gabija anti-phage mechanism

Liu-DS1 was annotated as a Gabija-like system in our analysis. Canonical Gabija systems encode two proteins: GajA, which contains an ATPase-like domain and a TOPRIM (topoisomerase-primase) nuclease domain, and GajB, a UvrD-family helicase (*18, 34–36*). GajA and GajB form a supramolecular complex that executes cell death to inhibit phage replication (*34, 36*). In contrast, the *Pseudomonas aeruginosa* Gabija has been proposed to directly sense and cleave phage DNA ends, resulting in protection of the infected cells (*37*). Compared to typical Gabija, Liu-DS1 contains a GajB homologue and a hypothetical protein instead of GajA (hereafter GajC) (Fig. 2A). The Liu-DS1 GajB contains the conserved GajB UvrD helicase domain, together with an additional domain of unknown function (Fig. S4A).

**Fig. 2.**
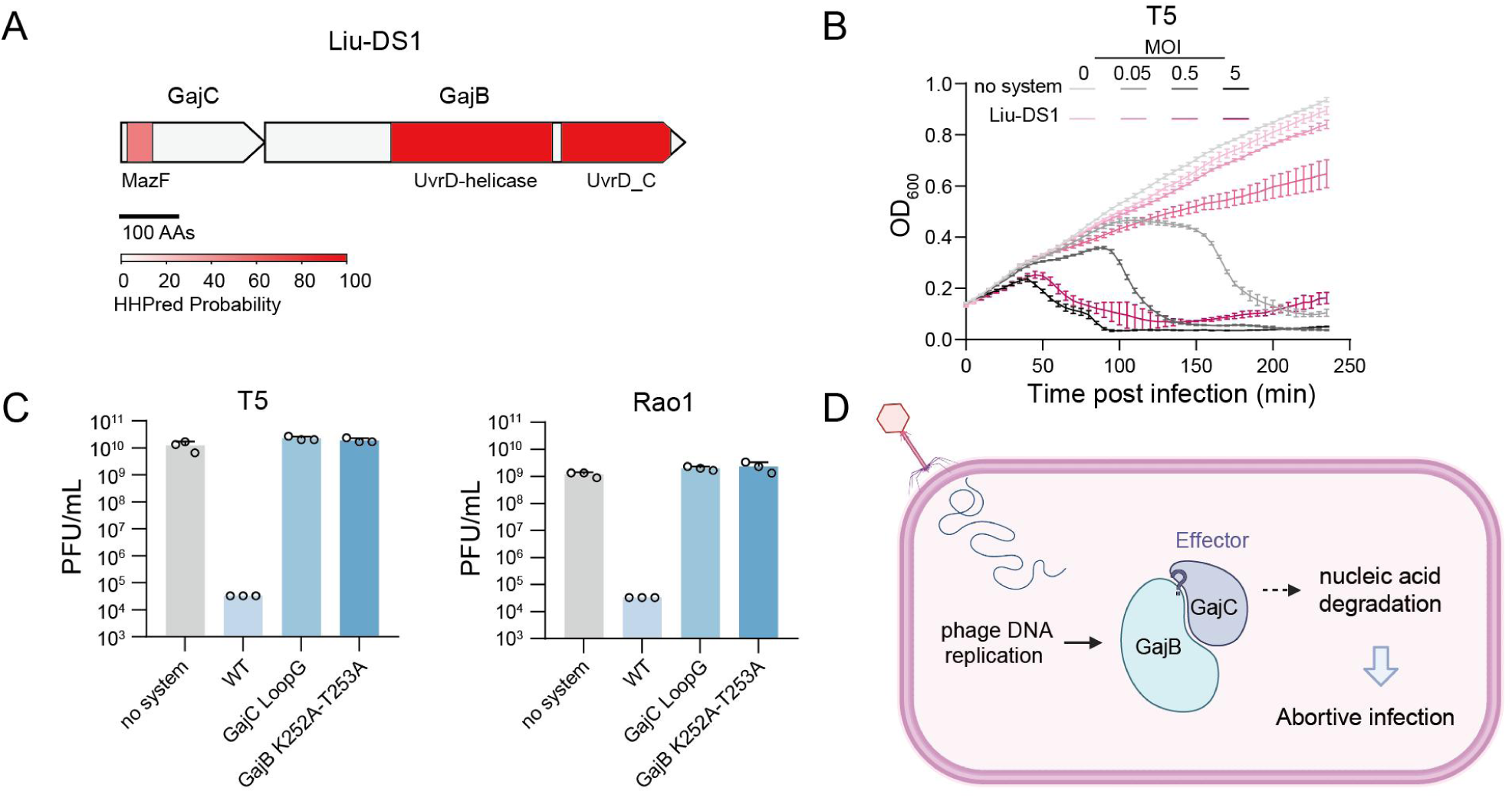
Liu-DS1 represents a Gabija subtype. (A) Domain organization of Liu-DS1. (B) Growth curves of *E. coli* expressing Liu-DS1 and no system after infection with T5 at MOIs of 0.05, 0.5 and 5. Data represent the mean ± SD of n = 3 biological replicates. (C) Plaque forming units (PFUs) of T5 and Rao1 infecting *E. coli* strains expressing wild type (WT) or mutated Liu-DS1 systems. LoopG: substitution of β1-β2 loop with glycine residues. Data represent the mean ± standard deviation of n = 3 biological replicates, with individual data points overlaid. (D) Predicted model for anti-phage defense by Liu-DS1. GajB is hypothesized to recognize phage infection by sensing DNA ends or ssDNA and activate GajC for nucleic acid degradation, resulting in cell death or dormancy and phage replication suppression. Created with BioRender.com.

Structure prediction of GajC reveals that it adopts a split MazF-like domain interrupted by a 5 α-helix insertion (Fig. S4B). MazF is a well-characterized RNA endonuclease from the MazEF toxin-antitoxin system that induces cell death through RNA degradation (*38*). The structural similarity between GajC and MazF suggests that GajC acts as the nuclease effector in Liu-DS1.

Plaque forming assays showed that Liu-DS1 provides immunity against 20 of the tested 71 phages (Fig. 1C). Analysis of growth curves of the liquid cultures expressing Liu-DS1 post infection indicated that Liu-DS1 only protected bacterial culture at a low multiplicity of infection (MOI < 1), whereas infection at MOI of 5 resulted in culture collapse (Fig. 2B), indicating that Liu-DS1 provides population-level immunity via triggering an abortive infection (Abi) response (*7*). To probe the underlying mechanism, we analyzed mutated variants of the system.

Mutation of the Walker A motif in GajB (K252A-T253A) abolished defense, in line with the necessity of the helicase domain in Gabija systems (*34, 36*)(Fig. 2C).

Structural comparison of GajC and MazF revealed that GajC features a β1-β2 loop corresponding to the active site of MazF (*38, 39*). Substitution of this loop with glycine residues abolished immunity (Fig. 2C). These results suggest that both GajB and GajC activities are required for the immune function of Liu-DS1. However, because GajC lacks an ATPase-like domain, it is unlikely to be able to sense ATP depletion as GajA (*34–36*). We speculate that GajC might be activated by its direct interaction with GajB, which may detect atypical DNA structures such as phage DNA ends or replication intermediates, similar to the *P. aeruginosa* Gabija and other nuclease-helicase systems (*37, 40, 41*) (Fig. 2D).

### CBASS recruits a diadenylate cyclase to trigger immune signaling

Liu-DS2 is annotated as a CBASS system because it contains a cGAS-like cyclase and a Patatin-like phospholipase (Fig. 3A), a typical gene architecture of the type I CBASS system (*25*), in which the cyclase produces an immune signal upon sensing infection, which activates the phospholipase effector to degrade the cell membrane and trigger cell death. In addition to these canonical components, Liu-DS2 encodes a second enzyme containing a diadenylate cyclase domain (DAC) (Fig. 3A). Notably, Liu-DS2 was identified in a recent preprint but provided only preliminarily characterization (*42*). Our analyses reveal that Liu-DS2 provided protection against 5 phages and protected cultures only at a low MOI, consistent with an Abi response (Fig. 3B).

**Fig. 3.**
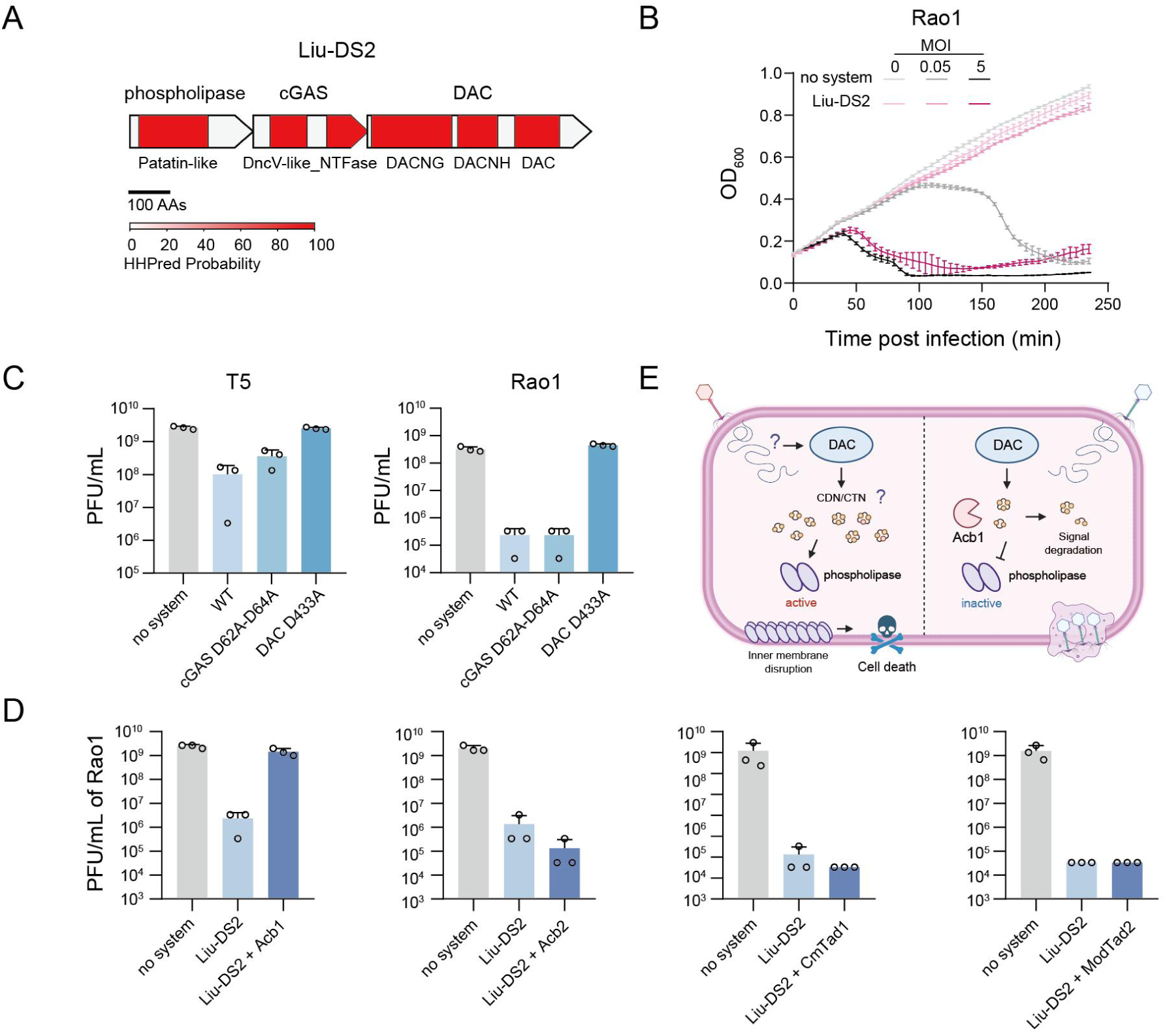
Liu-DS2 is a CBASS subtype with a diadenylate cyclase for triggering immune signaling. (A) Domain organization of Liu-DS2. (B) Growth curves of *E. coli* expressing Liu-DS2 and no system after infection with Rao1 at MOIs of 0.05 and 5. Data represent the mean ± SD of n = 3 biological replicates. (C) Plaque forming units (PFUs) of T5 and Rao1 infecting *E. coli* strains expressing wild type (WT) or mutated Liu-DS2 systems. Data represent the mean ± standard deviation of n = 3 biological replicates, with individual data points overlaid. (D) Effects on the indicated anti-defense proteins on the PFUs of Rao1 infecting *E. coli* expressing Liu-DS2. Data represent the mean ± standard deviation of n = 3 biological replicates, with individual data points overlaid. (E) The proposed model for anti-phage defense by Liu-DS2 and the anti-Liu-DS2 function of Acb1. Left: DAC senses phage infection through unknown mechanism and produces a cyclic-nucleotide signal, which activates the Patatin effector to destruct membrane integrity. Right: Acb1 degrades the cyclic-nucleotide signal and inhibits the Liu-DS2 immunity. Created with BioRender.com.

To dissect the function of DAC in Liu-DS2, we constructed mutated systems in which the conserved catalytic residues of DAC and cGAS were replaced with alanine, respectively. Disruption of DAC abolished the immunity against T5 and Rao1, whereas the cGAS mutation exerted minor effects (Fig. 3C), indicating that the enzymatic activity of DAC, rather than cGAS, is essential for defense. To investigate whether Liu-DS2 acts through a nucleotide-based signaling molecule, we analyzed the effects of anti-defense proteins that degrade or sequester the nucleotide signaling molecules. The results show that Acb1, a phosphodiesterase that cleaves adenosine-containing cyclic di-or trinucleotide signals (*43*), efficiently suppressed Liu-DS2 immunity, indicating that the immunity relies on a nucleotide-based signaling molecule (Fig. 3D). In contrast, the sponge proteins Tad1, Tad2 and Acb2, which have been shown capable of sequestering a wide range of immune signals, could not inhibit Liu-DS2 (*44, 45*) (Fig. 3D). Overall, our data support a model in which Liu-DS2 derives from the recruitment of a diadenylate cyclase into a CBASS operon, forming a hybrid signaling system in which the DAC synthesizes the immune signal that activates the downstream phospholipase effector (Fig. 3E).

### Acquisition of putative helicases by DarTG and ietAS systems

Liu-DS3 was annotated as DarTG system, a toxin-antitoxin system that provides phage defense via an Abi response (*46*). Canonical DarTG systems contain two components: DarT, a DNA ADP-ribosyltransferase toxin that modifies phage DNA and prevents phage replication, and DarG, the antitoxin protein that neutralizes the activity of DarT (*46*). In contrast, Liu-DS3 encodes an additional predicted helicase, hereafter TgaH (Dar<u>TG</u>-<u>a</u>ssociated <u>h</u>elicase) (Fig. 4A). Structural prediction showed that TgaH resembles UvrD-family helicases (DALI Z score: 27.1) but lacks the 2B domain (Fig. S5A), a region implicated in dsDNA binding but not essential for the helicase activity (*47, 48*).

**Fig. 4.**
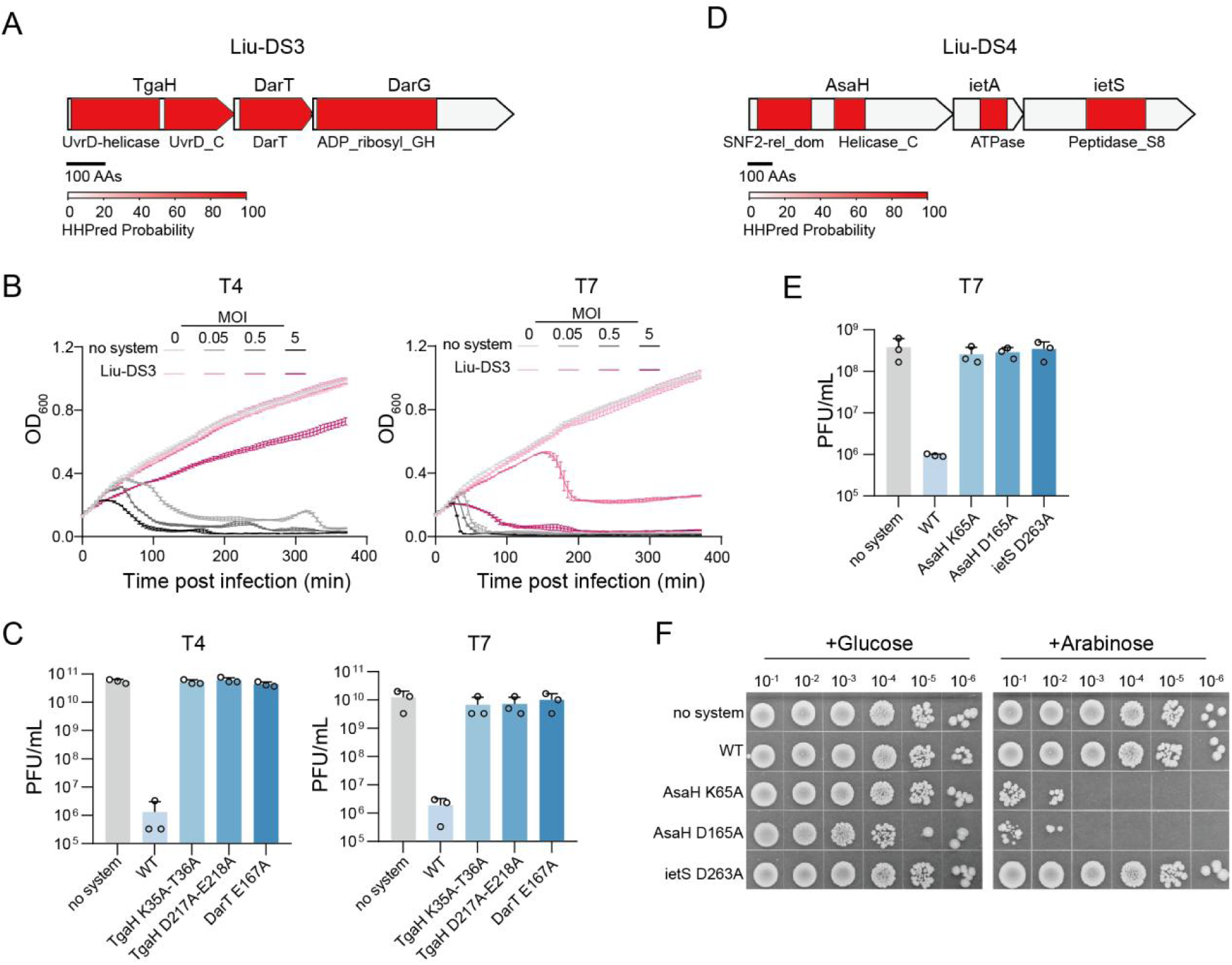
DarTG and ietAS systems employ helicase-like proteins as essential components. (A) Domain organization of Liu-DS3. (B) Growth curves of *E. coli* expressing Liu-DS3 and no system after infection with T4 and T7 at MOIs of 0.05, 0.5 and 5. Data represent the mean ± SD of n = 3 biological replicates. (C) Plaque forming units (PFUs) of T4 and T7 infecting *E. coli* strains expressing wild type (WT) or mutated Liu-DS3 systems. Data represent the mean ± standard deviation of n = 3 biological replicates, with individual data points overlaid. (D) Domain organization of Liu-DS4. (E) PFUs of T7 infecting *E. coli* strains expressing wild type (WT) or mutated Liu-DS4 systems. Data represent the mean ± standard deviation of n = 3 biological replicates, with individual data points overlaid. (F) The AsaH mutation induced cytotoxicity. Strains containing empty vector or expressing WT or mutated Liu-DS4 were cultured in medium containing glucose (to repress protein expression) or arabinose (to induce protein expression), and then the cell viability was analyzed by plotting on agar plates.

Our phage challenge assays revealed that Liu-DS3 resulted in culture collapse post T7 infection at high MOI, but allowed retarded culture growth post infection of T4, T5, and SSL_2009a at an MOI of 5 (Fig. 4B, Fig. S6). These phenotypes are consistent with an Abi response but suggest that DarT toxicity may be regulated in this system, enabling partial growth under certain infection conditions. Moreover, Liu-DS3 must be activated sufficiently early to inhibit the killing of host cells by the replication of T4, T5, and SSL_2009a. Mutation of the DarT active site or the conserved ATP-binding residues of TgaH abolished the anti-phage function, indicating that TgaH indeed plays an essential role in immunity (Fig. 4C).

Liu-DS4 is an ietAS system with an additional helicase, hereafter AsaH (iet<u>AS</u>-<u>a</u>ssociated <u>h</u>elicase) (Fig. 4D). IetAS was originally identified as a plasmid stabilization toxin-antitoxin system, encoding a serine protease as the toxin and an ATPase as the antitoxin (*49*). However, the anti-phage mechanism of ietAS remains unknown (*19*). In Liu-DS4, the AsaH helicase belongs to the SNF2 family and resembles the DrmD helicase of the Class I DISARM system (Fig. 4D, Fig. S5B) (*50*). Mutation of the conserved Walker A motif (K65A) and the DEAH box (D165A) of AsaH abrogated the protection (Fig. 4E), indicative of the essential role of AsaH in anti-phage function. The protease activity is also critical for the anti-phage function, in line with previous studies (*19*). Our attempts to construct a plasmid expressing the system carrying a mutated ietA were unsuccessful, reinforcing that the active ietA functions as the antitoxin (*49*). Moreover, induction of Liu-DS4 carrying mutated AsaH resulted in cytotoxicity (Fig. 4F), suggesting that AsaH also contributes to antitoxin-like regulation, analogous to toxin-antitoxin-chaperone systems (*51, 52*).

Together, we reveal that the putative helicases recruited by DarTG and ietAS systems are indeed required for their anti-phage functions.

### Synergistic immunity of Pycsar and Thoeris

Liu-DS6 is a five-gene operon (Fig. 5A), annotated as a Pycsar system in our analysis. The first 2 genes, encoding a transmembrane (TM) protein (DS6-1, PycTM) and a cyclase (DS6-2, PycC), represent a typical Pycsar system (*53*). The third and fourth genes constitute a type IV Thoeris system, containing a TIR-domain protein and a Caspase-like protein (*54*), and the fifth gene encodes a protein of unknown function. Both Pycsar and Thoeris synthesize nucleotide-based signaling molecules, which then activate their cognate effectors to trigger an immune response (*53, 55*).

**Fig. 5.**
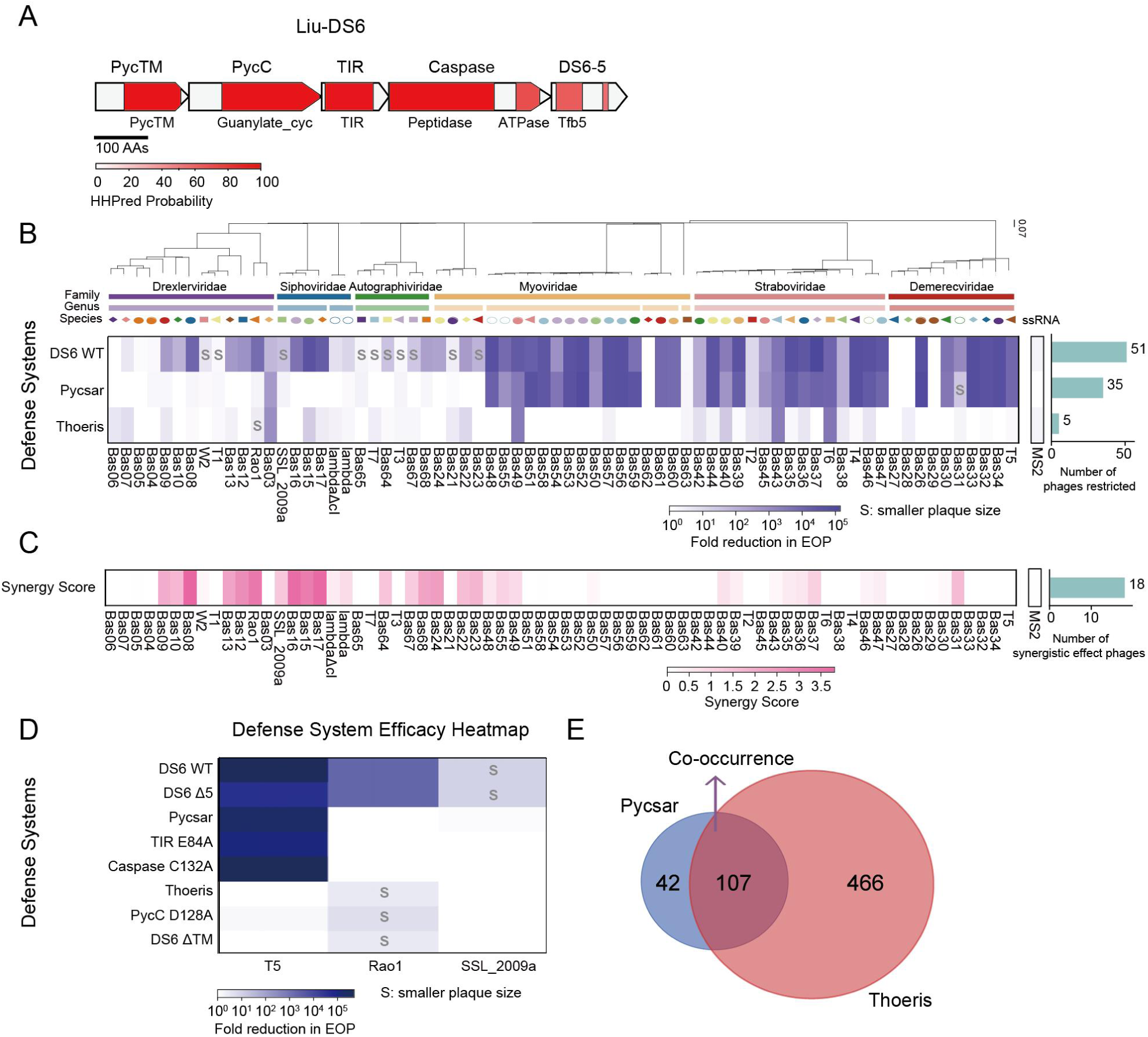
Type IV Thoeris and Clade F Pycsar exhibit synergistic anti-phage activity. (A) Domain organization of Liu-DS6. (B) Defense heatmap of the anti-phage activity of full-length Liu-DS6, and Pycsar and Thoeris alone, showing decrease in efficiency of plating (EOP) by blue squares and reduced plaque size indicated by the letter “s”. The decrease fold of EOP was calculated by 3 independent experiments. The number of restricted phages, the EOP reduction fold of which was more than 101.5, were counted. (C) Synergy score of the anti-phage activity, calculated by comparing the immunity of full-length Liu-DS6 with the sum of Pycsar and Thoeris immunity. The number of phages with synergy score >1 was counted. (D) Defense heatmap of the anti-phage activity of full-length Liu-DS6, Pycsar and Thoeris alone, and mutated Liu-DS6 against T5, Rao1 and SSL_2009a. The decrease fold of EOP was calculated by 3 independent experiments. (E) Numbers of strains encoding either DS6 Pycsar homologous systems (42), or Type IV Thoeris systems (466), or both systems in one operon (107).

To examine the phylogenetic relationship between Liu-DS6 Pycsar and previously characterized Pycsar systems, we searched for the homologous systems of DS6 Pycsar by PSI-BLAST with both DS6-1 and DS6-2 sequences, and then constructed a phylogenetic tree using the resulting DS6-2 homologues and the 967 PycC proteins identified previously (*53*). Most DS6-2 homologues clustered into a distinct branch that is distantly related to known PycC proteins, with a minority scattered in Clade C and E of PycC, both of which employ TM effectors. These results indicate that Liu-DS6 encodes a previously unrecognized clade of Pycsar (designated Clade F) (Fig. S7). Notably, earlier surveys focused on cNMP cyclases in defense islands (*53*), which likely contributed to clade F being overlooked.

We then aimed to investigate whether the Pycsar and Thoeris components of Liu-DS6 function independently. To this end, we constructed strains either expressing Pycsar from the native promoter or Thoeris under an arabinose-inducible promoter.

Challenging the strains with our collection of phages showed that both Pycsar and Thoeris independently defend against several phages (Fig. 5B). However, the combined set of phages restricted by either Pycsar or Thoeris individually was smaller than the set restricted by the full-length Liu-DS6 operon, suggesting synergistic activity. Calculation of the epistatic coefficients, comparing the immunity of the full operon with the sum of Pycsar and Thoeris individually, confirmed widespread synergy against a wide range of phages (Fig. 5C).

To investigate whether the synergy is mediated by cross-activation of the TM and Caspase effectors, we constructed chimeric systems expressing TIR and PycTM, and PycC and the Caspase-like protein, respectively. The chimeric systems were challenged with T5 which is only sensitive to Pycsar, and SSL_2009a, which is only sensitive to Thoeris. This revealed that the chimeric systems did not provide protection (Fig. S8), indicating the effectors are not cross-activated.

We next examined which components of Liu-DS6 are required for the synergy. We constructed mutated Liu-DS6 systems that lack the 5^th^ gene (Liu-DS6^Δ5^) or PycTM (Liu-DS6^ΔTM^), or carry mutations within the active sites of PycC, TIR, and Caspase. Liu-DS6^Δ5^ showed similar immunity to the whole operon against the tested phages, indicating that the 5^th^ gene plays a minor role under our tested conditions (Fig. 5D). In contrast, deletion of PycTM and mutation of PycC, TIR, and Caspase eliminated the synergistic effect, resulting in the same immune activity as the remaining intact system. The data suggest that full activity of both Pycsar and Thoeris is essential for the synergistic effect (Fig. 5D).

Given the synergistic effect of the Pycsar and Thoeris systems, we investigated how often the two systems co-occur in one operon. Our analysis detected 149 Pycsar homologue systems and 573 type IV Thoeris systems across ∼46,000 prokaryotic genomes, with 107 of them co-occurring in one operon (Fig. 5E, Table S6). These hybrid operons are exclusively present in Gammaproteobacteria (Fig. S9) and 90 of the 107 operons contain the 5^th^ gene (Table S6). The prevalence of Pycsar-Thoeris combinations suggests that their synergistic defense is biologically advantageous, and the possible accessory role of the fifth gene warrants further investigation.

### Prevalence and distribution of the characterized defense systems

To understand how widespread these newly identified systems are, we analyzed their prevalence and phylogenetic distribution. We first assessed their occurrence in *E. coli* (Fig. S10A). Liu-DS1 (GajC-GajB) and Liu-DS4 (AsaH-ietAS) showed the highest abundance, encoded by more than 1.5% of *E. coli* genomes, whereas the other 4 systems are encoded by less than 0.5% of genomes. We then analyzed their distribution across bacterial phyla (Fig. S10B-G). Notably, Liu-DS2 (CBASS-DAC), Liu-DS4, and Liu-DS6 (combined Pycsar and type IV Thoeris) are exclusively encoded by *Pseudomonadota*, likely reflecting a sampling bias towards *Pseudomonadota*, since these systems were identified from *E. coli*. Nevertheless, two systems were more frequently found in other bacterial phyla: Liu-DS3 (TgaH-DarTG) was most abundant in *Thermodesulfobacteriota* (∼3.6%) and *Bacteroidota* (∼2.6%), while Liu-DS1 is more often encoded in *Thermodesulfobacteriota* (∼0.7%). Together, these analyses show that the six validated systems are distributed across diverse bacterial lineages and expand the known diversity of the bacterial anti-phage repertoire.

## Discussion

The perpetual arms race with phages has driven the evolution of a widespread and diverse repertoire of anti-phage defense systems in bacteria. Diversification of these systems is often facilitated by their modular organization, which enables frequent recruitment and exchange of defensive modules (*2, 3, 21, 24–31*). In this study, we leveraged this modular nature to discover new defense systems by systematically searching for operons encoding defensive domains but exhibiting non-canonical architectures. After excluding known defense systems and potential housekeeping operons, the remaining operons represent candidate new defense systems. These candidate defense operons (CDOs) outnumbered complete defense systems detected by DefenseFinder by ∼2.2-fold in *E. coli* strains, suggesting that the abundance and diversity of anti-phage systems remain substantially underestimated, even in the most intensively studied bacterial species.

From the 9 selected CDOs, we validated 6 functional systems, including 2 systems (Liu-DS2, CBASS-DAC, and Liu-DS5, HamMAB) recently discovered independently by other groups during the preparation of this manuscript (*33, 42*). Overall, the validated systems can be classified into 3 categories: (1) acquisition of additional modules by known systems (Liu-DS2,3,4,5); (2) replacement of a defensive module with an alternative protein (Liu-DS1); (3) combination of independent but synergistic systems (Liu-DS6). Through functional characterization of these systems, we reveal that a variant of Gabija recruited a MazF-like domain to replace the canonical GajA nuclease, and that DarTG and ietAS, two toxin-antitoxin defense systems, have acquired in some instances helicases as essential components. The ietAS-associated helicase could control the ietA toxicity, similar to tripartite toxin-antitoxin systems that contain a molecular chaperone to stabilize the antitoxin module (*51, 52*). Our data further support that DAC in Liu-DS2 generates a nucleotide-based signal, which can be cleaved by Acb1 to prevent effector activation. Interestingly, cGAS is not required for the immunity against Rao1 or T5. A likely explanation is that cGAS and DAC may sense different phages, in analogy to Thoeris systems that employ multiple TIR proteins for recognition of different phage-associated molecular patterns (*55*).

Our approach also uncovered a new clade of Pycsar systems (Clade F), which was missed by previous screening of cNMP cyclases from defense islands (*53*). Clade F Pycsar shows a marked tendency to co-occur with type IV Thoeris in one operon, and we reveal that the two systems exhibit synergistic immunity against diverse phages. Although synergy has been observed between non-co-localized systems (*56*), organization within an operon may enable tighter coordination and facilitate simultaneous horizontal gene transfer. Such organization would provide a selective advantage to the host and promote the spread of the operon during the arms race with phages, consistent with the prevalence of combined Thoeris-Pycsar operons in *Pseudomonadota*. A previous study revealed that synergistic systems may co-opt sensor modules to enhance anti-phage activity (*56*). However, our results show no detectable cross-talk between the sensor and effector modules of the chimeric systems, suggesting that the signal-effector interactions are highly specific. The molecular basis for the synergistic effect of Thoeris-Pycsar requires further investigation.

The guilt-by-association principle has been proven a powerful method to predict new defense systems (*18, 19*), but systems that typically reside outside of defense islands or nearby unknown defense systems (*11*) can be potentially missed. Although our strategy incorporates a guilt-by-association filter, it uses a broader criterion (proximity of a known defense system within 50 kb) than previous studies (*18, 19*). This less restrictive framework allowed us to discover systems that are not frequently located in defense islands, as exemplified by Clade F Pycsar. Together, our findings highlight the power of a defense domain-centric strategy to uncover novel bacterial immunity architectures and reveal new routes for anti-phage system diversification. We expect that applying this framework more broadly beyond *E. coli* will continue to expand the bacterial anti-phage defense repertoire.

## Materials and Methods

### Prediction of candidate defense operons

A total of 4,243 high-quality *E. coli* genomes that retain GCF identifiers were downloaded from NCBI and analyzed by DefenseFinder (*8*) (v1.0.9, hmmer 3.3). This generated two files, protein_defense_finder_genes.tsv and protein_defense_finder_hmmer.tsv, which listed proteins in complete defense systems (Table S1) and proteins with the HMM profile of a protein from known defense systems respectively. The proteins in the latter but absent from the former (n = 551,238) were defined as defensive domain-containing proteins and extracted for further analysis.

To detect operons encoding defensive domain-containing proteins, the five upstream and five downstream genes were extracted from the genome annotation file (GFF). Of the 11 genes (including the defensive domain-encoding gene), those on the same coding strand with intergenic gaps ≤100 bp were considered an operon. One-gene operons were removed. Repeated operons with 100% identity were removed, generating 214,164 unique operons. These operons were automatically annotated by the names designated by DefenseFinder (Table S2).

To exclude potential housekeeping operons, 50 kb upstream and 50 kb downstream of the genomic context from the defensive domain-encoding gene were analyzed by DefenseFinder. Only when at least one known defense system was detected in the 100 kb region, the defensive domain-encoding gene was considered as a seed. Finally, candidate defense operons (CDOs) were detected with the seeds as described above (Table S3). To further remove redundancy, these CDOs were clustered with CD-HIT(-c 0.9-n 5) (v4.8.1) (*57*), resulting in 3,919 representative CDOs (Table S4).

### Strains and phages

Plasmid construction was performed in *E. coli* DH5α. Phage assays were performed with *E. coli* MG1655 as hosts. DH5α and MG1655 were cultured at 37℃ with shaking at 180 rpm in standard LB broth; strains used for phage assays were cultivated under same conditions in MMB medium (LB supplemented with 0.1 mM MnCl_2_ and 5 mM MgCl_2_). For plasmid maintenance, media were fortified with streptomycin (50 μg ml^-1^) or ampicillin (100 μg ml^-1^) as appropriate. All phages used in this study, together with their corresponding hosts, are listed in Table S9. Rao1 phage was kindly provided by Prof. Shuke Wu (Huazhong Agricultural University). Phages of the BASEL collection were a generous gift from Dr. Alexander Harms (University of Basel).

### Plasmid construction

The plasmid vectors used in this study and the constructed plasmids are listed in Table S7. To express the candidate systems, the coding sequences together with their native promoters indicated in Table S5 were synthesized by General Bio (Anhui, China) and then cloned into the pBADHisA vector. For Liu-DS5 and DS7-9, initial plaque assay using the systems with their own native promoters did not detect anti-phage activity. Therefore, the coding sequences of these systems were also inserted into pBADHisA vector between *EcoR*I and *Nco*I restriction sites, such that expression of the genes can be driven by an araBAD promoter. Site-directed mutagenesis was carried out by PCR amplification of plasmids containing the target genes with mutagenic primers, and the resulting fragments were assembled in *E. coli* DH5α.

To co-express the anti-defense proteins Acb1, Acb2 with Liu-DS2, pCDFaras vector was constructed by replacing the T7 promoters of pCDFDuet-1 with araBAD promoter from pBADHisA. Acb1 and Acb2 were then amplified from T4 phage and inserted into pCDFaras. Genes encoding CmTad1 and ModTad2 were synthesized by General Bio (Anhui, China) and cloned into pCDFtet, in which the T7 promoters of pCDFDuet-1 were replaced by the tetR/tetA promoter.

To express the components of Liu-DS6 in two plasmids, truncated Liu-DS6 operon was still inserted into pBADHisA, where the co-expressed genes were amplified from the complete system and inserted into the pCDFtet vector between the *Pco*I and *Xho*I.

All the plasmids were constructed using ClonExpress II One Step Cloning Kit (Vazyme, C112-02), and verified by Sanger sequencing (Wuhan Tianyi Huayu GeneTechnology, China) or through whole-plasmid sequencing with Nanopore technology (Jiangsu COWIN BIOTECH, China). The primers used for cloning are summarized in Table S8.

### Phage Propagation

To propagate phages, an overnight culture of *E. coli* MG1655 was diluted 1:100 into fresh MMB broth (LB plus 0.1 mM MnCl₂ and 5 mM MgCl₂) and shaken at 37 °C until the absorbance of 600 nm (OD_600_) reached 0.3. 5 mL of this culture was then inoculated with the desired phage at an MOI of 0.01 and further incubated for 4 h.

Cellular debris was removed by centrifugation (5000 × g, 10 min) and the clarified lysate was passed through a 0.22 µm syringe filter (Millipore SLGPM33RS). The sterile phage stock was stored at 4 °C.

To determine the phage titer, a double-layer agar method was employed. Serial ten-fold dilutions of each phage lysate were prepared in SM buffer (50 mM Tris–HCl pH 7.5, 100 mM NaCl, 8 mM MgSO_4_). For each dilution (10^-3^ to 10^-8^), 3 µL was spotted onto double-layer plates prepared by mixing 200 µL of an overnight MG1655 culture with 10 mL molten MMB soft agar (0.5 % w/v) and immediately overlaying onto MMB bottom agar (1.5 % w/v). After overnight incubation at 37 °C, plaques were counted and the titer was calculated.

### Plaque assay

*E. coli* MG1655 cells carrying the indicated plasmids were subjected to plaque formation assays. The cells were grown overnight at 37 °C in MMB medium supplemented with the corresponding antibiotics. A 200 µL aliquot of the culture was mixed with 10 mL of molten MMB soft agar (0.5%) containing the same antibiotics and overlaid onto MMB plates (1.5%) supplemented likewise. To induce protein expression from the araBAD promoter, 0.2% (w/v) L-arabinose was generally added, except Acb1, which was induced with 0.05% (w/v) L-arabinose. For CmTad1 and ModTad2, 10 ng/mL anhydrotetracycline (aTc) was added to the mixture as the inducer. Ten-fold serial dilutions of the tested phage in SM buffer were then spotted (3 µL each) onto the bacterial lawn, followed by 8 h of incubation at 37°C before imaging.

### Phage infection assays in liquid culture

Overnight cultures of *E. coli* MG1655 carrying either empty pBAD-HisA or expressing a system were diluted 1:100 into fresh MMB medium containing ampicillin (100 µg·ml^-1^) and grown at 37 °C with shaking until OD_600_ reached 0.2. 180 µL of each culture were then dispensed into 96-well microplates. Phage suspensions were prepared in SM buffer and 20 µL aliquots were added to final multiplicities of infection (MOIs) of 0.05, 0.5 or 5; control wells were supplemented with 20 µL SM buffer (MOI = 0). Plates were incubated in a FLUOstar OMEGA microplate reader (BMG Labtech, Germany) at 37 °C with orbital shaking at 200 rpm, and the OD_600_ was recorded every 5 min for the duration of the experiment.

### Cell toxicity assay

The function of AsaH in antitoxin activity was assessed by cell toxicity assay. Overnight cultures of *E. coli* MG1655 harboring the empty vector pBAD-HisA or plasmids encoding wild-type or mutated Liu-DS4 systems were diluted 1:100 in MMB medium supplemented with ampicillin (100 µg·mL^-1^) and grown at 37°C to an OD_600_ of ∼0.2. The cultures were then induced with 0.2% arabinose or suppressed with 2% glucose and incubated for 3 hours at 37°C. Subsequently, 10-fold serial dilutions (from 10^-1^ to 10^-6^) were prepared in MMB, and 5 µL of each dilution was spotted onto MMB plates containing ampicillin (100 µg·mL^-1^). After overnight incubation at 37 °C, plates were imaged and colonies were counted.

### Distribution analysis of the validated systems

A total of 46,026 complete bacterial genomes were downloaded from RefSeq database and analyzed by DefenseFinder to annotate defensive domain-containing proteins. To detect the homologous systems of the DS6 Pycsar and Thoeris, homologues of PycC and Ths-Caspase were detected by the corresponding HMM profiles (Pycsar AG_cyclase.hmm and Thoeris_IV ThsA_Casp4.hmm, respectively). Operons encoding these proteins were obtained as described in the section “Prediction of candidate defense operons” and proteins from the resulting operons were pooled. The protein pool was searched by PSI-BLAST (v2.14.1+) with three iterations (e-value=1 × 10^-3^) using DS6 PycTM as the query to detect operons encoding both a DS6 PycTM homologue and PycC. These operons were considered to contain homologous systems of DS6 Pycsar. Type IV Thoeris systems were detected with the same procedure using DS6 TIR as query for PSI-BLAST. This pipeline was also used to detect co-occurrence by searching for ThsCaspase homologues in the protein pool of operons encoding DS6 Pycsar, and to analyze the association of DS6-5 with DS6 Pycsar and Thoeris. The resulting detected systems were manually inspected for false positives (Table S6). A similar pipeline was employed to analyze the distribution of other validated systems.

### Phylogenetic tree construction

To construct the phage phylogenetic tree, phages in this study were clustered using the Genome-BLAST Distance Phylogeny (GBDP) method with the D0 formula for nucleotides in VICTOR (*58*). Taxonomy data were obtained from the NCBI database or the BASEL phage collection (*59*).

To construct a phylogenetic tree of organisms encoding DS6 Pycsar and/or Thoeris, 16S rRNA sequences from the corresponding genomes were extracted using RNAmmer. Multiple sequence alignment was performed with MAFFT (v7.526) (*60*) under --auto parameters, followed by trimming with trimAl (v1.4.rev15)(*61*) using default settings. IQ-TREE (v2.2.6) (*62*) executed a two-step modeling process: first identifying the optimal substitution model via-m MF option, then constructing the final phylogenetic tree with 1,000 ultrafast bootstrap replicates (-bb 1000) with the SYM+I+R7 model.

To generate the phylogenetic tree in Fig. S6, protein sequences, including 35 non-redundant DS6 PycC sequences, 967 previously identified PycC sequences and 8 outgroup sequences, were aligned according to the procedure described above with the LG+R8 model, and the tree was generated with the iTOL online tool (*63*).

### Domain annotation and structure prediction

The proteins of Liu-DS1-6 were analyzed by HHpred with default parameters (*64*) to identify functional domains. Structures of these proteins were predicted by AlphaFold3 (*65*), and DALI search server was used to identify homologous structures (*66*). The predicted structures and superimposed structures were visualized by PyMOL 3.0.3.

## Statistics Analysis

For all statistics analysis, the assays were performed in three independent replicates as indicated in the figure legends.

## Code availability

Codes used in this study can be found at https://github.com/yanqiuLiu0908/Identification-of-Defense-Associated-Operons.

## Acknowledgments

We thank Fangkui Wang at the National Key Laboratory of Agricultural Microbiology Core Facility and Jinshan Li at Experimental Teaching Center of Bioengineering, Huazhong Agricultural University for technical support. We thank Prof. Shuke Wu for providing Rao1 phage and Dr. Alexander Harms for providing BASEL collection.

## Funding

National Key Research and Development program of China 2022YFA0912200 (W.H.); National Natural Science Foundation of China 31970545 (W.H.); National Natural Science Foundation of China 32270099 (W.H.); Fundamental Research Funds for Central Universities 2662024SKPY003 (W.H.); Hubei Special Project for Science Development 2024CSA060 (W.H.).

## Author contributions

Conceptualization: W.H., Y.L. Methodology: Y.L., W.H., R.P-R.

Investigation: Y.L., K.T., Z.H., Y.C., K.X., P.L., R.F., F.L.

Visualization: Y.L., K.T. Supervision: W.H.

Writing—original draft: W.H., Y.L. K.T., R.P-R. Writing—review & editing: All authors

## Competing interests

The authors declare no competing interests.

## Data and materials availability

All data are available in the main text or the supplementary materials.Codes used in this study can be found at https://github.com/yanqiuLiu0908/Identification-of-Defense-Associated-Operons.

## Supplementary Materials

Figs. S1 to S10

Tables S1 to S9

---

**Fig. S1.**
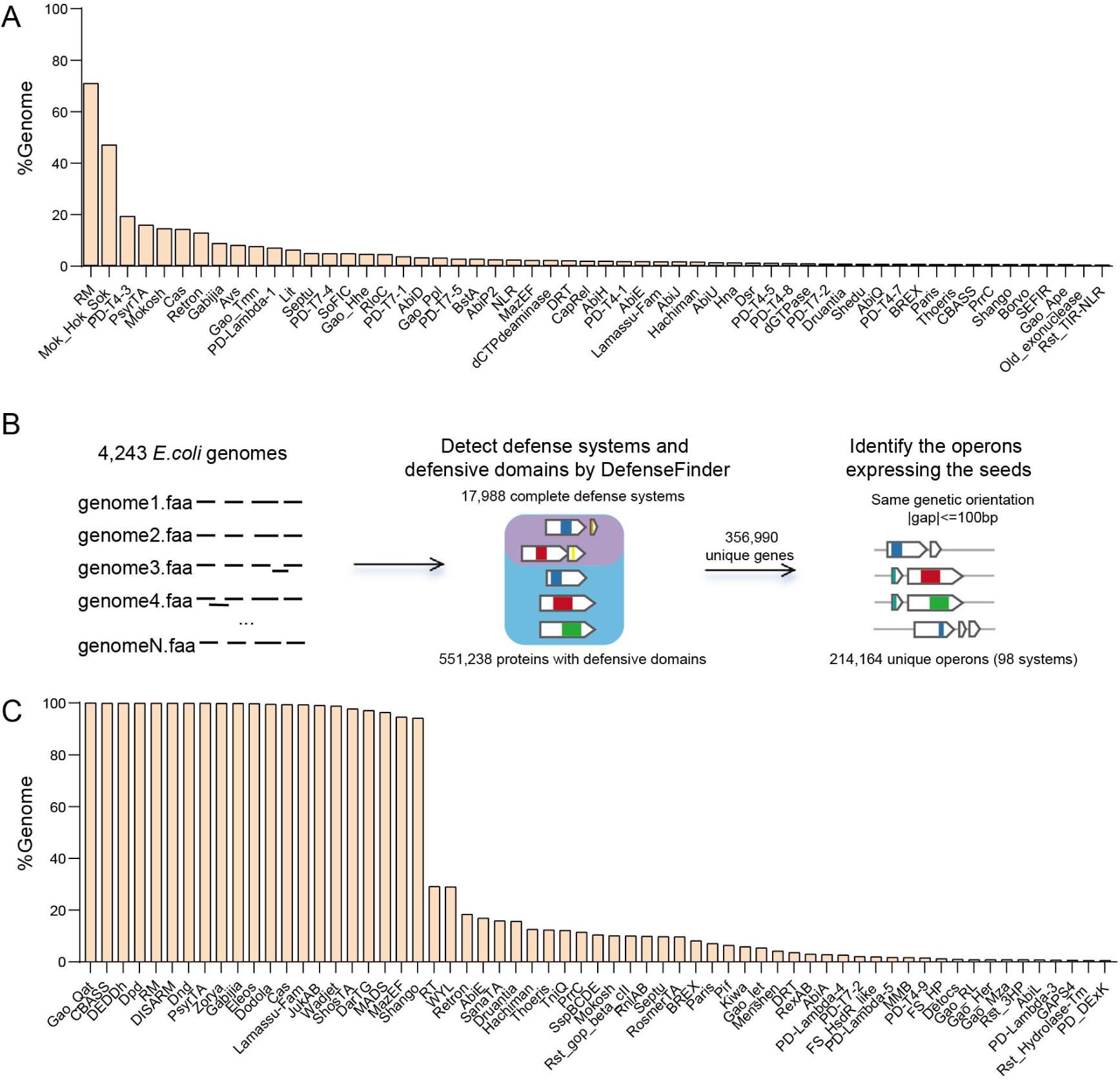
(A) Frequency of complete defense systems in *E. coli* genomes. (B) Strategy used for the identification of operons encoding defensive domains in *E. coli* genomes. (C) Frequency of operons encoding defensive domains in *E. coli* genomes. The frequency of 21 operon types is almost 100%, indicating that the strategy shown in (B) is used to detect housekeeping operons.

**Fig. S2.**
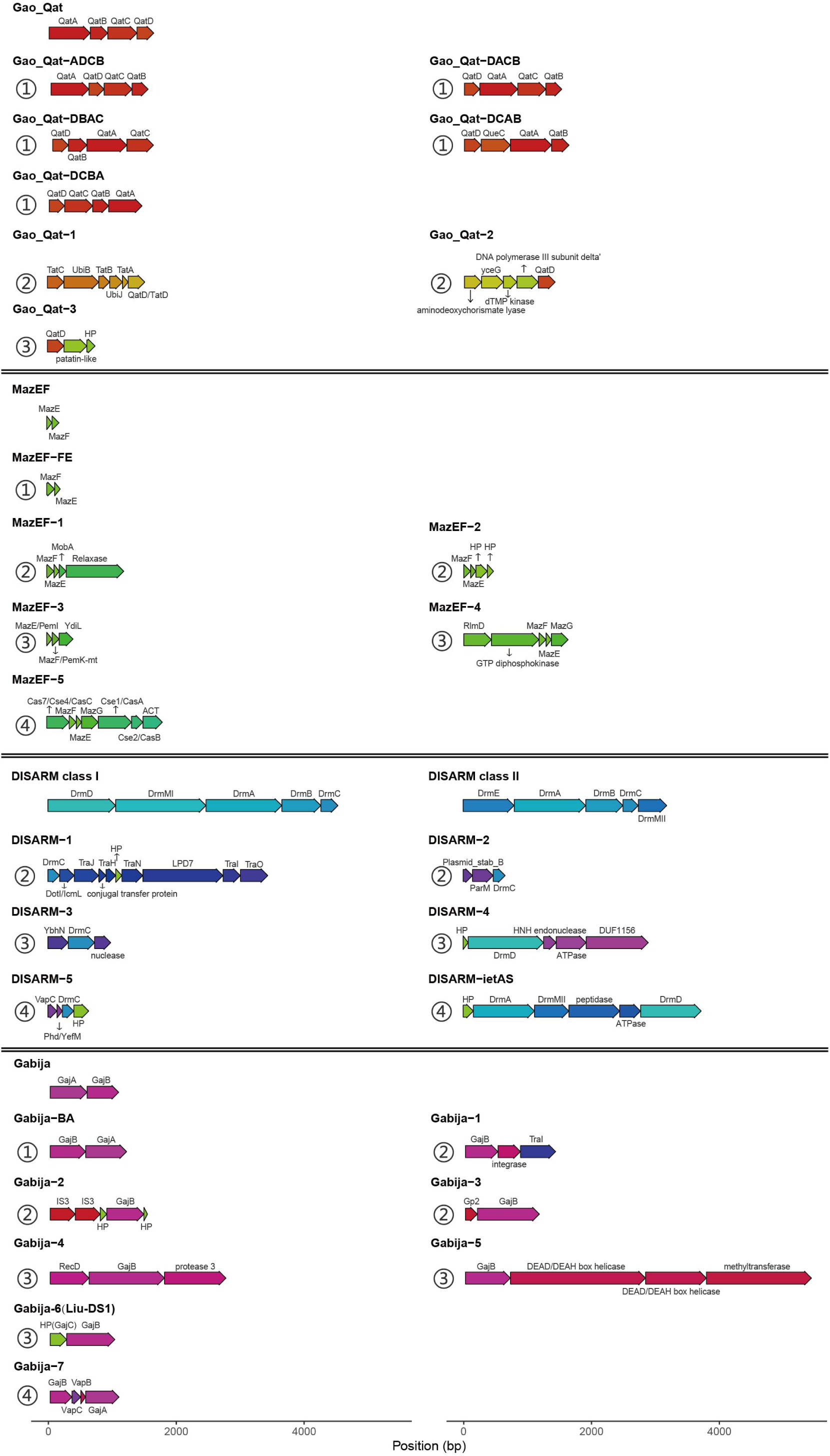
Operon architectures of known Gao-Qat, MazEF, DISARM and Gabija defense systems and the corresponding candidate defense operons (CDOs). The CDOs identified in our analysis can be classified into 4 types: (1) complete defense system with a different gene order, i.e., Gao_Qat-ADCB, (2) association of a defensive component with non-defensive proteins, i.e., Gao_Qat-1, (3) new combination of defense components from different systems, i.e., Gao_Qat-3, (4) association of defense system with another defense system, i.e., Gabija-7.

**Fig. S3.**
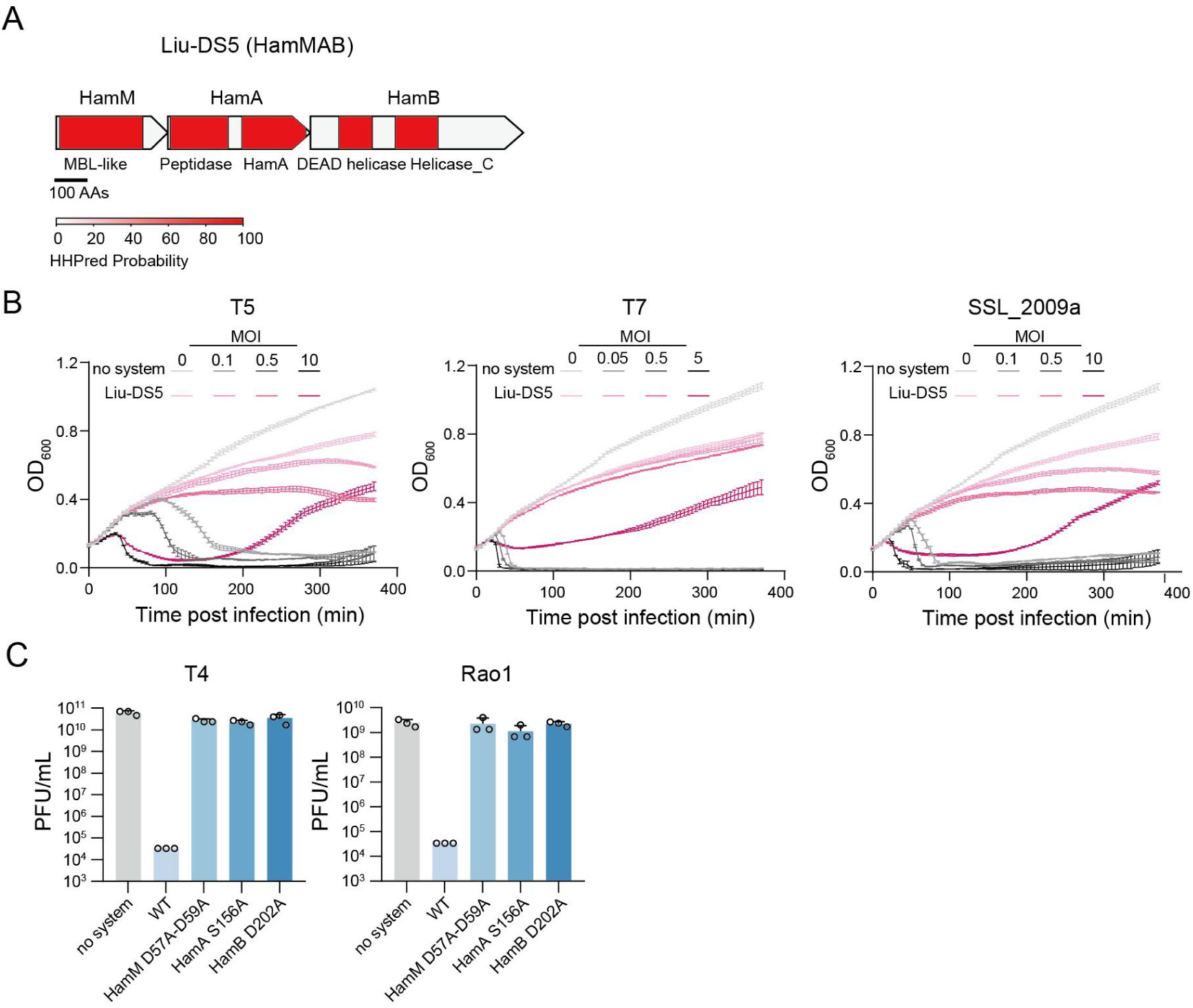
Liu-DS5 (HamMAB) is a fused system consisting of Hachiman and a nuclease-protease module. (A) Domain organization of Liu-DS5. (B) Growth curves of *E.coli* expressing Liu-DS5 and no system after infection with T5 and SSL_2009a at MOIs of 0.1, 0.5 and 10, and with T7 at MOIs of 0.05, 0.5 and 5. Data represent the mean ± SD of n = 3 biological replicates. (C) PFUs of T4 and Rao1 infecting *E. coli* strains expressing wild type (WT) or mutated Liu-DS5 systems. Data represent the mean ± standard deviation of n = 3 biological replicates, with individual data points overlaid.

**Fig. S4.**
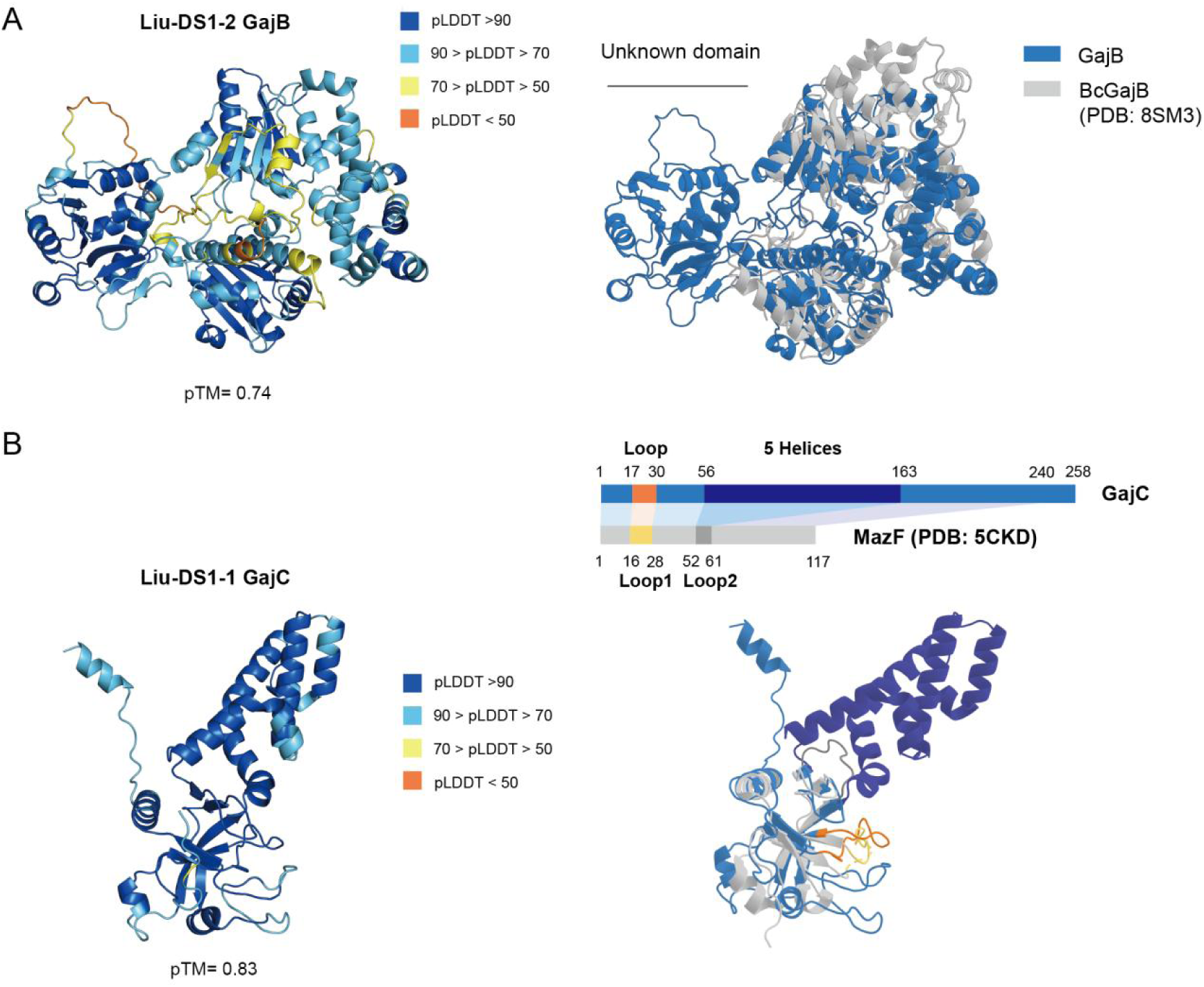
Structural prediction of GajB and GajC. (A) Left: predicted structure of Liu-DS1-2 GajB by AlphaFold 3 (pTM = 0.74), colored by predicted local distance difference test (pLDDT). Right: Structural comparison of Liu-DS1-2 GajB (blue) and *Bacillus. cereus* VD045 GajB (grey, PDB ID 8SM3) with a root mean square deviation (RMSD) of 6.762 Å over 136 equivalent residues. (B) Left: AlphaFold3-predicted structure of Liu-DS1-1 GajC (pTM = 0.83) and colored by pLDDT. Right: Structural comparison of Liu-DS1-1 GajC (blue) and MazF (grey, PDB ID 5CKD) with an RMSD value of 1.875 Å for 241 Cα atoms, highlighting the key loop of GajC (orange) and loop1 of MazF (yellow) participating in catalytic activity. A schematic of the structural alignment of GajC and MazF is shown above the panel.

**Fig. S5.**
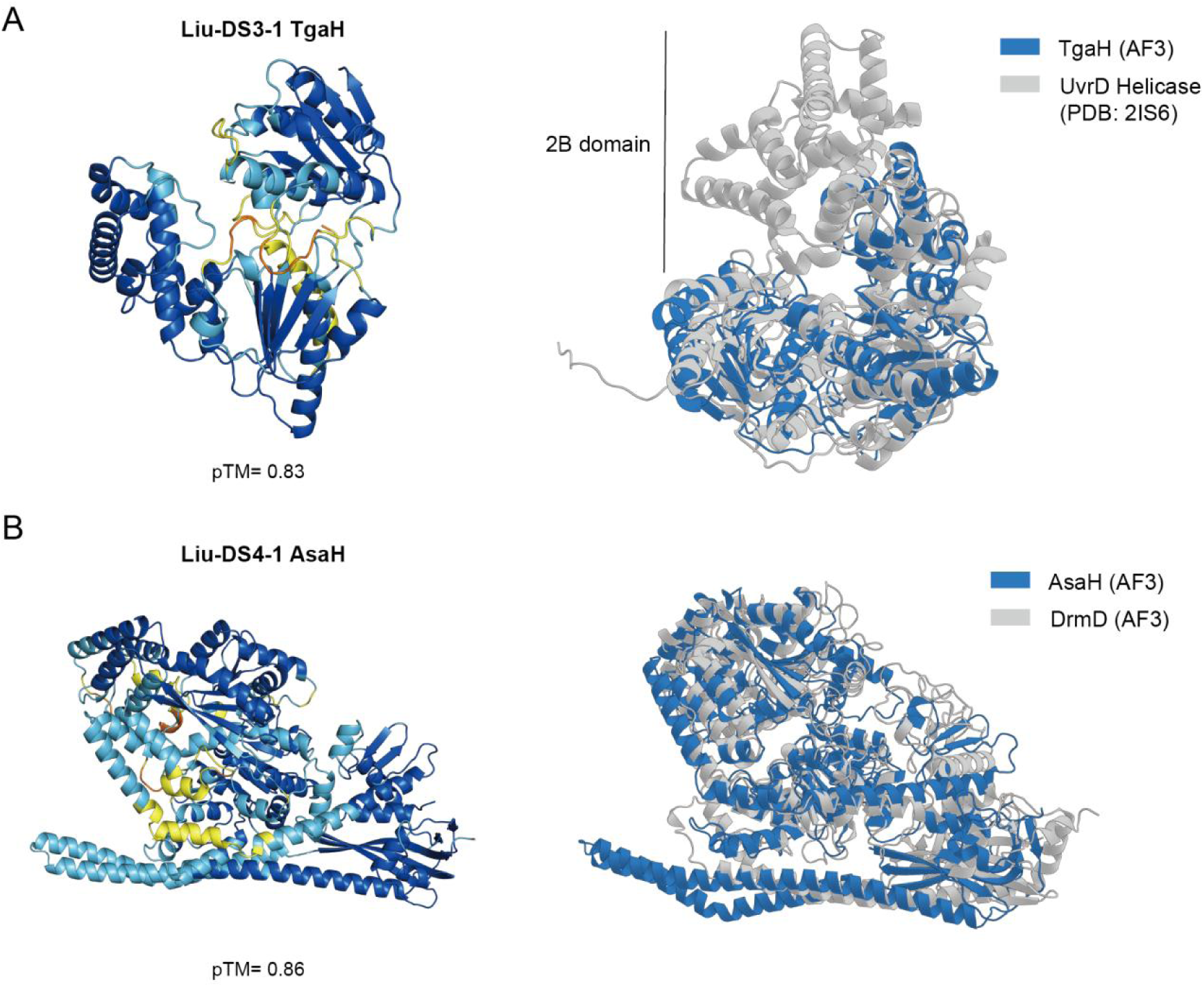
Structural prediction of TgaH and AsaH. (A) Left: predicted structure of TgaH by AlphaFold 3 (pTM = 0.83), colored by predicted local distance difference test (pLDDT). Right: Structural comparison of TgaH (blue) and a UvrD helicase (grey, PDB ID 2IS6) with a root mean square deviation (RMSD) of 5.119 Å over 296 equivalent residues. The 2B domain of UvrD is indicated. (B) Left: AlphaFold3-predicted structure of AsaH (pTM = 0.86) and colored by pLDDT. Right: Structural comparison of AsaH (pTM = 0.86) (blue) and AF3-predicted DrmD (grey, pTM = 0.9, WP_043276582.1) with an RMSD value of 3.758 Å over 312 equivalent residues.

**Fig. S6.**
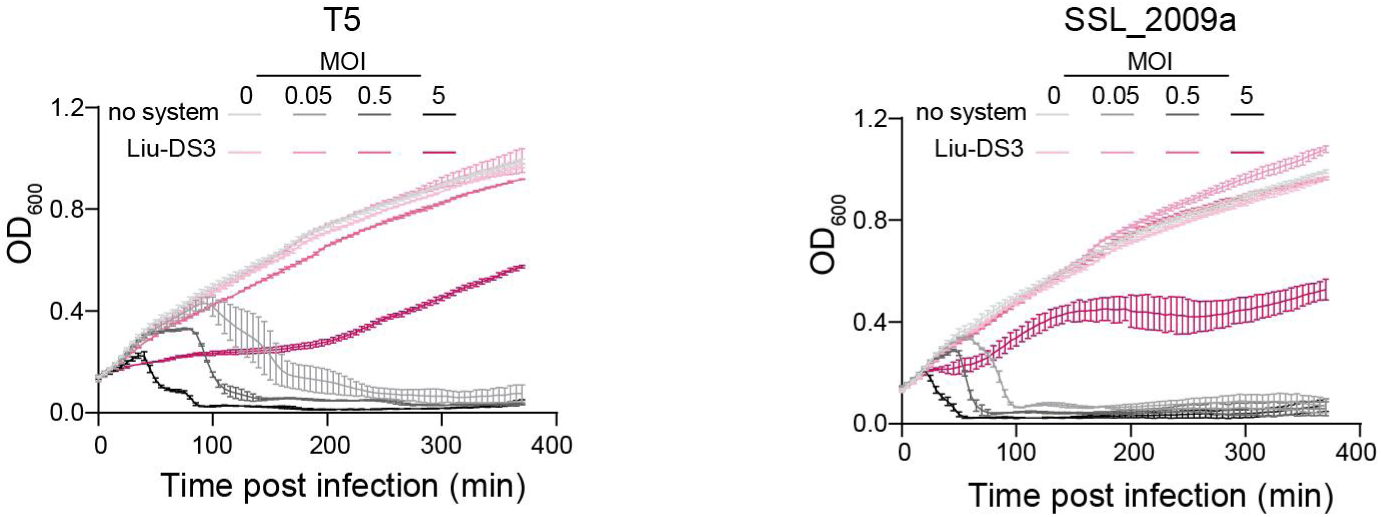
Growth curves of *E. coli* expressing Liu-DS3 and no system after infection with T5 and SSL_2009a at MOIs of 0.05, 0.5 and 5. Data represent the mean ± SD of n = 3 biological replicates.

**Fig. S7.**
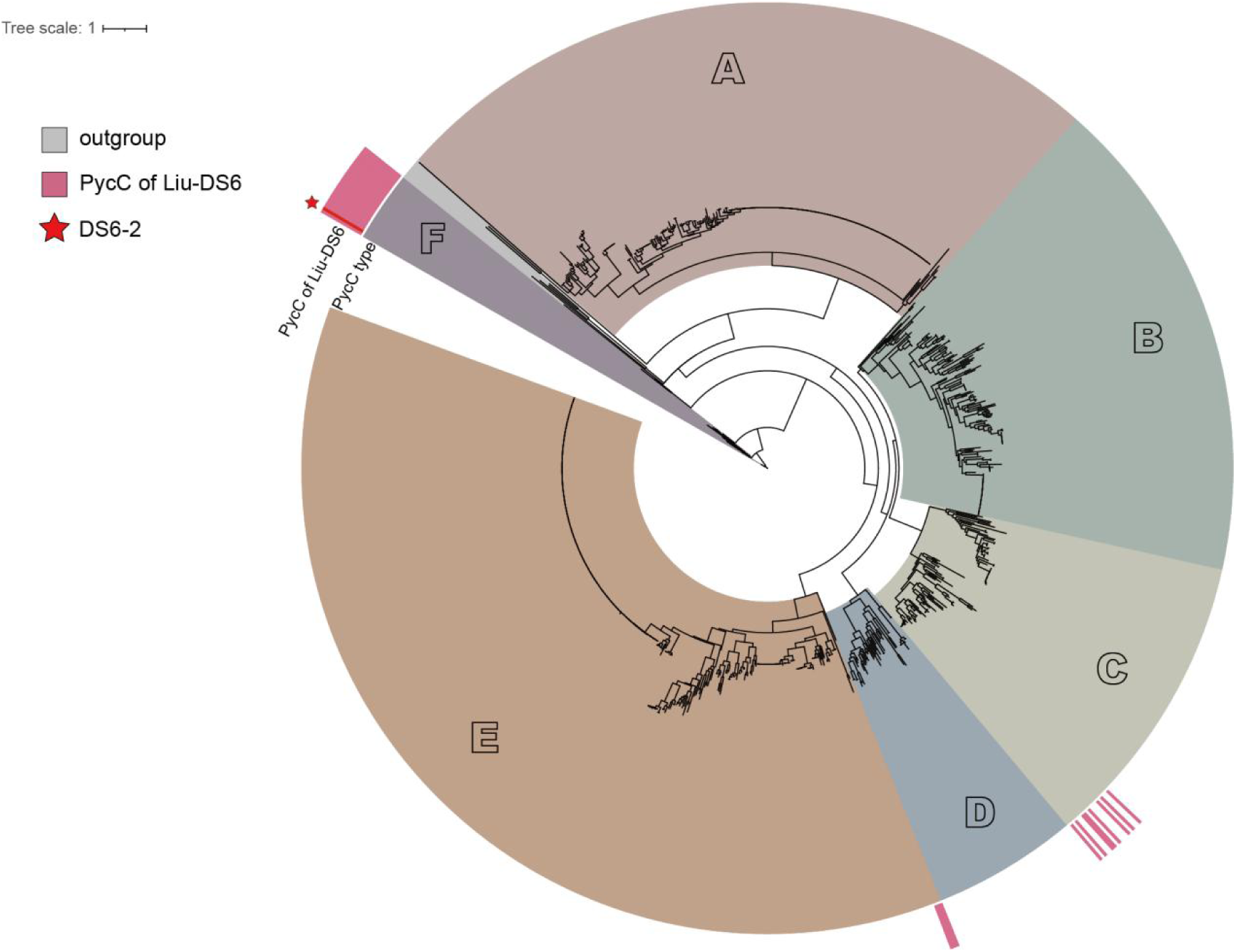
Phylogenetic analysis of PycCs from the DS6 homologous systems (highlighted by red bars) and previously identified PycCs reveals a novel PycC clade (F). Liu-DS6-2 is indicated with a red star.

**Fig. S8.**
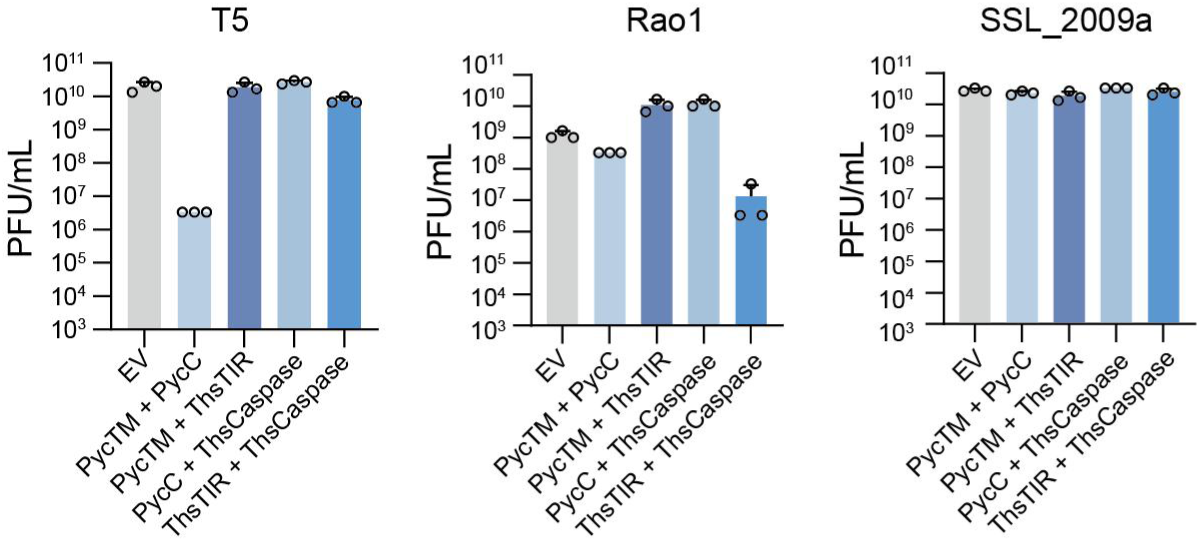
Chimeric systems expressing TIR and PycTM, or PycC and Caspase, show no immune activity against T5, Rao1 or SSL_2009a.

**Fig. S9.**
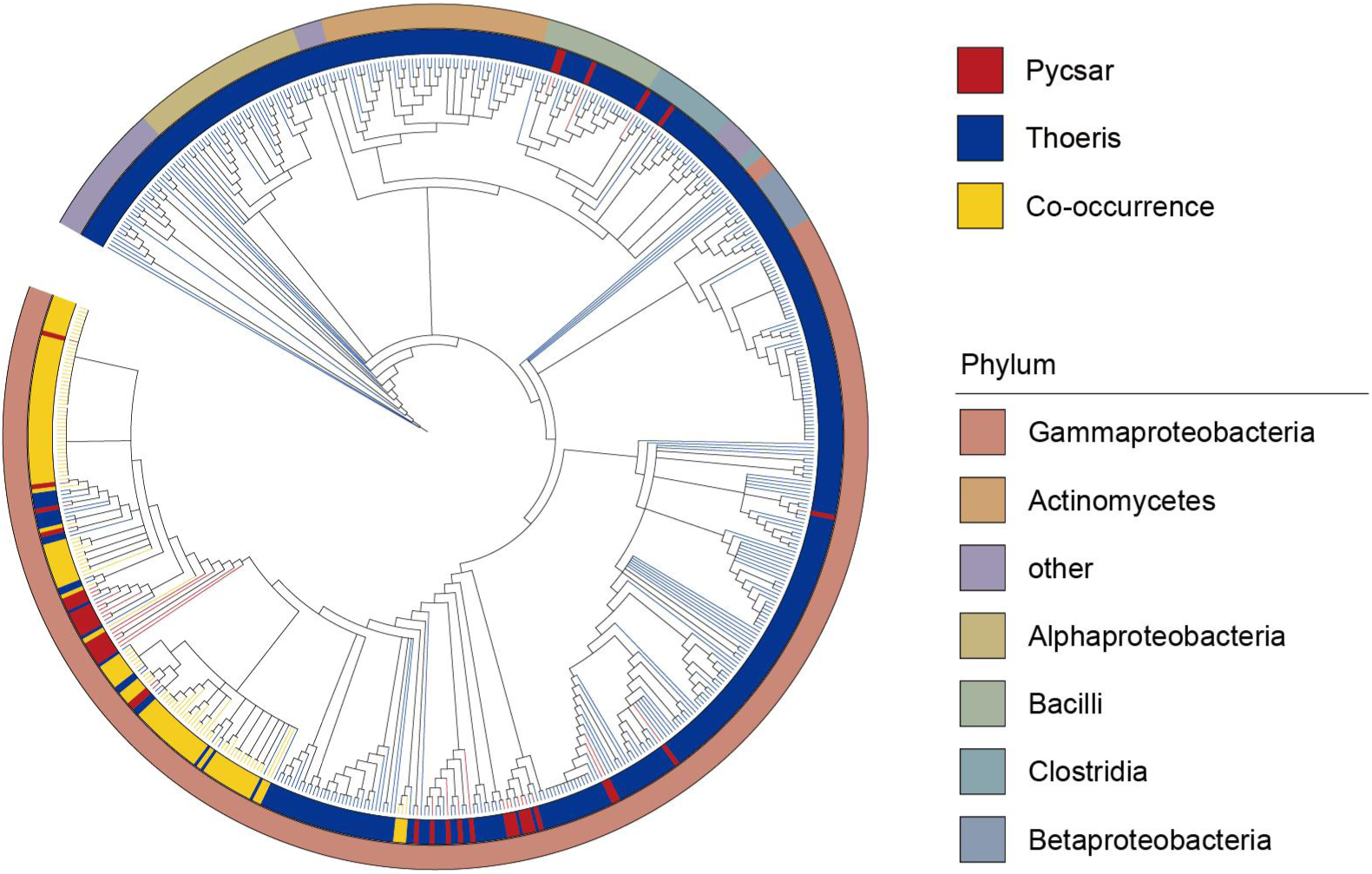
A phylogenetic tree of organisms encoding either DS6 Pycsar or Thoeris homologous systems, or both systems within one operon (co-occurrence). The outer ring indicates the bacterial phylum. Colors of the inner cycle indicate the presence of Pycsar (red), Thoeris (blue) or co-occurrence (yellow).

**Fig. S10.**
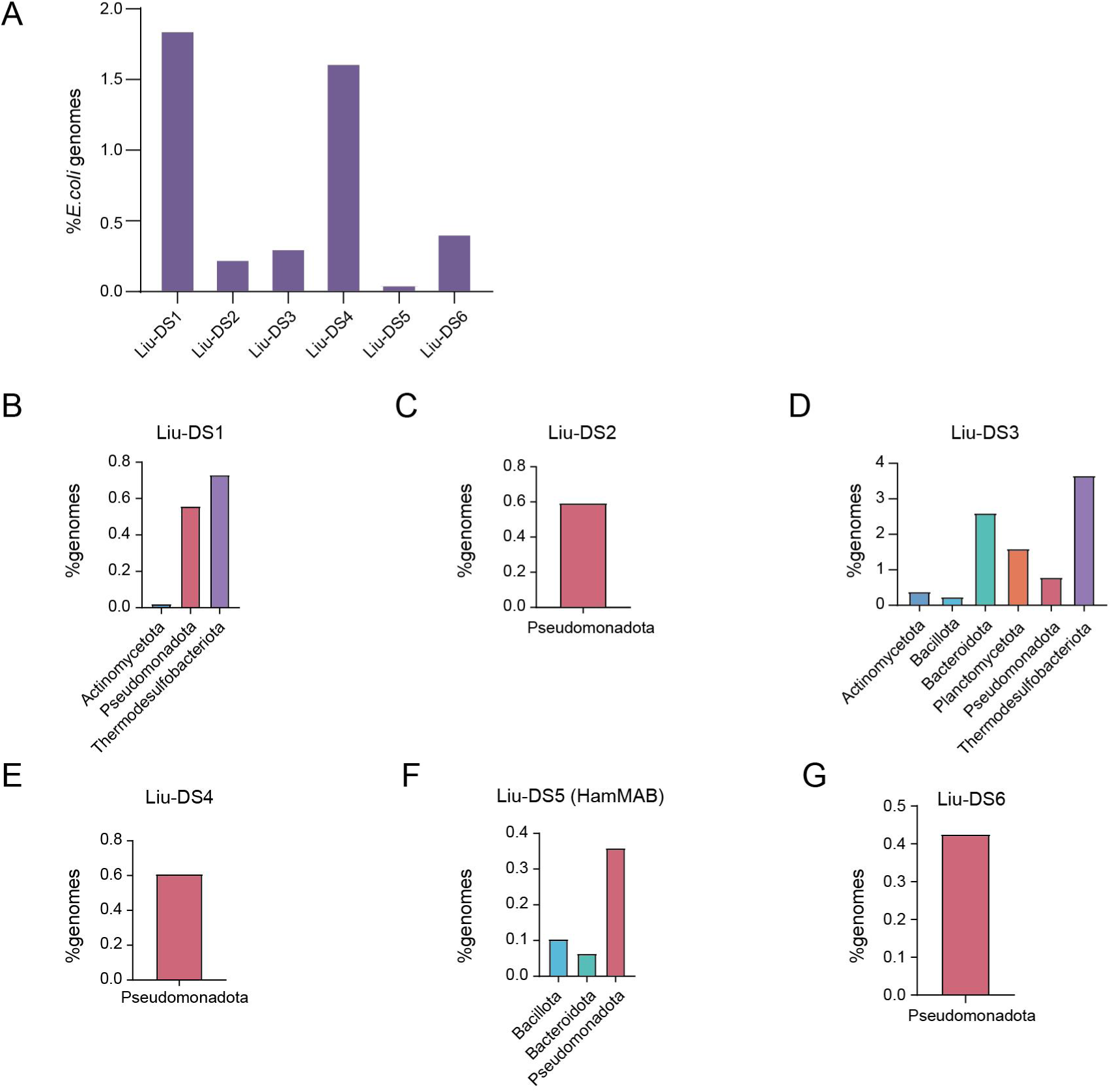
Abundance and distribution of the validated systems in this study. (A) The frequency of Liu-DS1-6 systems in 4243 *E. coli* strains. (B)-(G) The distribution of Liu-DS1-6 in bacterial phyla.

Table S1. Types and distributions of 17,988 complete defense systems in *E. coli* strains.

Table S2. Types and distributions of 214,164 operons that encode at least one defensive domain. The annotation of the operons was based on the names of known defense systems, which typically encode the defensive domain, and on the names of the HMM domains built into DefenseFinder.

Table S3. Types and distribution of 39,848 candidate defense operons.

Table S4. A list of 3,919 representative CDOs.

Table S5. Tested systems. The contig accession, source strains, encoded proteins and protein sequences are listed.

Table S6. A list of identified DS6 Pycsar or Thoeris homologous systems, or both systems within one operon (co-occurrence). Annotations of the proteins in the detected operons are shown.

Table S7. Plasmids used in the study. Table S8. Primers used in the study. Table S9. Phages used in this study.

## Notes

### Competing Interest Statement

The authors have declared no competing interest.

